# The role of TRPV4 in acute sleep deprivation-induced fear memory impairment

**DOI:** 10.1101/2024.08.12.607531

**Authors:** Meimei Guo, Feiyang Zhang, Sha Liu, Yi Zhang, Lesheng Wang, Jian Song, Wei Wei, Xiang Li

## Abstract

Acute sleep deprivation (ASD) negatively impacts fear memory, but the underlying mechanisms are not fully understood. Transient receptor potential vanilloid 4 (TRPV4), a cation channel which is closely correlated with the concentration of Ca^2+^, and neuronal Ca^2+^ overloading is a crucial inducement of learning and memory impairment. This study utilized an acute sleep-deprived mouse model combined with fear conditioning to investigate these mechanisms. mRNA sequencing revealed increased expression of TRPV4 in mice with ASD-induced fear memory impairment. Notably, knockdown of TRPV4 reversed ASD-induced fear memory impairment. ASD leads to the increased concentration of Ca^2+^. Additionally, we observed a reduction in spine density and a significant decrease in postsynaptic density protein 95 (PSD95), which is associated with synaptic plasticity, in sleep-deprived fear memory impairment mice. This indicates that ASD may cause overloaded Ca^2+^, disrupting synaptic plasticity and impairing fear memory. Moreover, TRPV4 knockdown significantly decreased Ca^2+^ concentration, mitigated the loss of dendritic spines and reduction of PSD95, contributing to the restoration of fear memory. These findings indicate a potential protective role of TRPV4 knockdown in counteracting ASD-induced fear memory deficits. Collectively, our results highlight that TRPV4 may be a potential therapeutic target in mediating fear memory impairment due to ASD and underscore the importance of sleep management for conditions like PTSD.

## Introduction

Sleep is crucial for normal life activities, regulating various brain physiological processes, including neuromodulation, neuronal activity, neurotransmission and others [1]. It plays an essential role in multiple cognitive processes [2, 3], particularly in supporting memory formation [4–6]. Sleep deprivation can lead to deficits in synaptic plasticity [7, 8], which is closely linked to learning and memory. Previous evidence shows that sleep deprivation impairs the consolidation of fear memory in both clinic and animal studies [6, 9, 10]. However, the precise mechanism of fear memory impairment caused by sleep deprivation remains unknown.

Fear memory is associated with the development and occurrence of trauma and emotional disorders, especially posttraumatic stress disorder (PTSD) [11]. The traditional and classical method to assess the expression of fear memory is fear conditioning, which relies on associative learning [12]. Noxious unconditioned stimuli (US) can alter the neuronal response to conditioned stimuli (CS), resulting in specific fear behaviors. Accumulating evidence suggests the significant role of long-term synaptic plasticity in the acquirement and encoding of fear memory [13, 14]. The medial prefrontal cortex (mPFC) is a crucial region related to the encoding and processing of fear memories [15–17]. The prelimbic (PL) region, a subdivision of the mPFC, is anatomically and functionally involved in fear behavior [18, 19]. PL cortices regulate stimulus-response and action-outcome learning [20] and play an essential role in the establishment of fear memories [21, 22]. For example, neuronal activities in the PL are closely related to freezing behaviors occurring during fear conditioning [21]. Stimulation of PL neurons contributes to enhanced freezing behaviors during the expression of fear memory [23].

Recently, it has been demonstrated that cation channels are associated with learning and memory [24, 25]. Transient receptor potential vanilloid (TRPV4) channels are broadly expressed in the brain and function as gated and non-selective cation channels, regulating many physiological processes [26]. Previous study suggested that the expression of TRPV4 is regulated by clock genes, which are linked with circadian rhythms [27, 28]. Disturbance of circadian rhythms significantly affects TRPV4 protein expression. Furthermore, TRPV4 can be activated by various stimuli, including circadian rhythm, temperature, and osmotic pressure, resulting in Ca2+ influx and inward currents [26, 27, 29]. Hyperactivation of TRPV4 channels, caused by exogenous and endogenous factors, enhances Ca2+ influx, leading to intracellular Ca2+ overload [30, 31]. Therefore, hyperactivated TRPV4 channels are involved in cognitive processes, such as hippocampal cell impairment induced by amyloid-β (Aβ) [32], and neuronal apoptosis [29]. Previous study have reported that the TRPV channel family is a risk factor for synaptic plasticity impairment [33] and neurobiological diseases, including Alzheimer’s disease [32, 34], brain edema [35, 36], cerebral ischemic reperfusion injury [37, 38]. However, it remains unclear whether TRPV4 is a vital factor for associative fear memory impairment induced by sleep deprivation, and how it alters following sleep deprivation and affects cued fear learning.

The postsynaptic density (PSD) is regarded as a dense area localized in excitatory synapses, including various receptors, signal molecules, and structural proteins related to synaptic plasticity. PSD95 is one of the most abundant proteins [39] involved in dendritic spine development, excitatory neurotransmitter transmission, and synaptic plasticity [40, 41]. Dysfunction of PSD95 is closely related to alterations in synaptic plasticity and impairment of learning and memory [42]. Knockout of the transient receptor potential cation channel 6 (TRPC6) has been shown to enhance the expression of PSD95 and alleviate learning and memory dysfunction by reducing Ca2+ influx [43].

Given these backgrounds, we hypothesize that TRPV4 affects sleep deprivation-induced fear memory impairment via reduced expression of PSD95 and synaptic plasticity damage. This study aims to characterize the role of TRPV4 in synaptic plasticity associated with fear memory impairment induced by sleep deprivation.

## Materials and methods

### Mice

Nine-week-old male C57BL/6J mice were given free access to food and water under a 12-hour light/dark cycle. All behavioral testing was conducted in red-light rooms. The protocols for this study were approved by the Animal Ethics Committee of Zhongnan Hospital of Wuhan University (Ethics approval number: ZN2023121).

### Acute sleep deprivation and fear conditioning test protocol

Before the experiment, the mice were allowed to acclimate to the environment and establish a circadian cycle (with the light phase from 8:00 am to 8:00 pm). On the experimental day, the mice were kept awake by gentle stimulation from 8:00 am to 2:00 pm [44]. During this period, animals were allowed to explore freely without disturbance until a sleep attempt was captured. At this point, they were kept awake using mild auditory and tactile stimulation using a pencil-sized paintbrush.

To minimize stress, the experimenter was forbidden to touch the mice at any time. After sleep deprivation, mice were allowed to stay at the operant chambers with a steel grid floor that can be energized for fear condition training. The context was scented with lemon flavor. Following a previous study’s protocol, the training included three trials to match the tone to the shock (80dB tone context stimulus (CS) for 120s and 0.6mA foot shock stimulus (US) lasting 1s) [22]. There were 120-second intervals between trials. The mice’s locomotion was captured by cameras in the chambers, and the data were processed and analyzed using software FreezeFrame 4 to acquire the percentage of freezing condition, which was defined as at least 1s of immobility status. The context control mice were exposed to the same context for 14 min, the same duration as the training protocol, without any tone or shock. The secondary day after training, the mice were returned to the lemon-scented context for recall tests. They were allowed to explore for 120 seconds, followed by three trails consisting of 120-second CS presentations alternating with 120-second intervals. The percentage of freezing time during the recall tests was assessed to evaluate memory retention.

### RNA sequence and data analysis

mRNA of mPFC tissue was extracted and rRNA was removed using kit from Vazyme (Nanjing, China). The Dynabeads purification kit (Invitrogen, USA) was then used to purify the mRNA. The mRNA was broken into fragments and cDNA libraries were constructed. Subsequently, the cDNA library was sequenced using Illumina HiSeq 4000 sequencing platform. The final data were mapped against the mouse genome (GRCm39) and read counts were analyzed to determine the expression of each gene across different groups (fold change > 1 and the *p* value < 0.05). Clustering analysis and Gene Ontology Enrichment Analysis were performed using the R packages.

### RNA extraction and RNA reverse transcription

According to the previous study [45], primarily, Trizol (Vazyme, Nanjing, China) was added in the ep tubes containing the tissues, which were then homogenized using electric grinding rods. Subsequent steps for RNA isolation were carried out as per the manufacturer’s instructions. The concentration of RNA was measured using a Nanondrop spectrophotometer (N50, IMPLEN, Germany). RNA reverse transcription was conducted using a kit from Vazyme (Nanjing, China) following the provided protocol. **Quantitative PCR**: After preparing the cDNA, quantitative PCR (qPCR) was performed following the manufacture’s protocol (Vazyme, Nanjing, China) in a 10ul reaction volume system (5ul SYBR green master mix, 2ul cDNA of each reaction, 500M primer per reaction). The concentration of cDNA was diluted according to the target gene abundance. All reactions were performed in duplicate using the Rotor-Gene Q PCR cycler. Data were normalized to the internal control PGK1 (phosphoglycerate kinase) and analyzed using the δδCT method. qPCR primers of each gene were listed below: TRPV4 F: AAACCTGCGTATGAAGTTCCAG; R: CCGTAGTCGAACAAGGAATCCA; Pim1 F: TGTCCAAGATCAACTCCCTGG; R: CCACCTGGTACTGCGACTC; ADNP F: GACTCCCACCACGAATCAGC; R: CCCGTTGAATTTAAGTTGGGCT; DAPK1 F: CAGATTCTCAGCGGCGTTTAC R: GATCCGAGGTTTGGGCACATT; PGK1 F: TGCACGCTTCAAAAGCGCACG; PGK1 R: AAGTCCACCCTCATCACGACCC.

### Western blot

PL tissue from mice were collected for protein extraction. The tissue was lysed using RIPA buffer (Servicebio, Wuhan, China) and denatured for 30 minutes at 65°C to ensure adequate cleavage of the protein. The concentration of protein samples was diluted to 5ug/ul with PBS and loading buffer. Proteins were separated by gel electrophoresis for 45 minutes and subsequently transferred onto a PVDF membrane (0000287894, Immobilon-P, Germany). The membrane was blocked using quick blocking buffer (HYC00811, HYCEZMBIO, Wuhan, China) for 30min at room temperature. Following blocking, the membrane was washed with Trizma-buffered saline containing 1% Triton-X-100 (TBST) lasting 7 minutes (three times). Then, the membrane was incubated with 10ml of rabbit monoclonal antibody (TRPV4 1:1000, A5660; PSD 95 1:1000, A7889; abclonal, Wuhan, China) in antibody diluent buffer for 18h at 4°C and washed using TBST for three times. Afterward, the membrane was incubated with HRP-rabbit anti-mouse antibody (1:10000, bioswamp, Wuhan, China) in antibody diluent for 1 hour and washed in TBST three times for 7min each time. The optical density and quantitative analysis of the blots were performed using the LI-COR analysis system.

### Immunofluorescence

The expression of PSD95 protein was measured in fear-training mice after sleep deprivation, following protocols from previous study [44, 46]. The sleep deprived mice were subjected to fear conditioning in operant chambers (Xinyi Instruments, Shanghai, China). Immediately afterward, the animals were anesthetized with isoflurane (RWD Life Science, Shenzhen, China) and perfused transcardially with PBS followed by paraformaldehyde. The brain tissues were collected, embedded in embedding medium, and sliced into 40 µm sections using a freezing microtome.

The brain slices were washed with PBST four times and incubated with blocking solution for 2h at 37 °C. Next, the brain slices were incubated with PSD95 antibodies (1:1000, Proteintech, 20665-1-AP, Wuhan, China) overnight at 4 °C. Following this, the slices were washed and incubated with goat anti-rabbit IgG conjugated to Alexa Fluor 594 (1:400, Jackson, USA) for 2 hours at 25 °C. After a final series of four washes, the slices were mounted on slides. Confocal microscopy (Leica, Germany) was used to capture the images, and ImageJ software was employed for further analysis.

### Lentivirus packaging

Lentivirus was packaged on the basis of the previous study [47]. HEK293T cells were used for the lentivirus production. When the cell density was grown to about 80% in single-layer flasks, the cells were transfected with three helper plasmids (pMDLg, pMDG and pRSV) and a transfer vector (constructed by cloning TRPV4 shRNA into the FG12 plasmid) using Lipofectamine 2000 transfection reagent, as per the manufacturer’s instructions. After 48 hours of culture, the supernatant was collected, filtrated and concentrated by ultracentrifugation. The concentrated virus, including one control lentivirus and another targeting TRPV4, were stored at −80 °C. The sequences for the shRNA were as follows: TRPV4 shRNA: CCCTGGCAAGAGTGAAATCTA; Universal control shRNA: GCGCGATAGCGCTAATAAT. The titers of each lentivirus were at least 1 x 10^8 IU/ml.

### Stereotaxic injection surgery

Mice were anesthetized with 1% pentobarbital (i.p., 50mg/kg) and placed in a stereotaxic frame (RWD Life Science, Shenzhen, China). During the experiment, a heating pad was used to keep animals warm. Control lentivirus and TRPV4 shRNA lentivirus (approximately 1 µL at a rate of 80 nL/min) were injected into the bilateral PL (2.34 mm anterior to the bregma, 0.20 mm lateral to the midline, and 1.90mm ventral from dura) using a glass pipette connected to a microinjection pump (RWD Life Science, Shenzhen, China). After injection, the pipette was remained in injection location for 10 minutes to prevent the spread of the virus. The brain wound was then sutured. Once the mice awoke, they were returned to their home cage and allowed at least one week to recover and ensure stable virus expression.

### Virus injection and analysis of dendritic spines

AAV (NCSP-RFP-2e5) was infused into the bilateral PL to reconstruct single neurons. After 21 days, sleep deprived and sleep ad libitum animals were subjected to fear conditioning in operant chambers. Immediately afterward, animals were anesthetized with isoflurane (RWD Life Science, Shenzhen, China) and perfused transcardially with PBS followed by paraformaldehyde. The brain tissues were collected, fixed in 4% PFA for 24-48 h, then sliced into 35 µm sections. The brain film slices were attached to an adhesive slide and sealed with anti-fluorescent quencher (Servicebio, Wuhan, China). The images were acquired by Zeiss 980 microscope with Airscan model. x63 oil immersion was used to capture the detail of dendritic spines. We selected the secondary basal dendrites by distance from the soma 50-110um as the previous study described [48] with z-stack model. Zeiss 980was used to further analyze the spine density in different groups. The experimenters were blinded to the groups and treatment in the whole course.

### Statistical analysis

The experimenter was blinded to the group assignments during the experiment. Data were analyzed using GraphPad Prism 9.0 (GraphPad, New York, USA), SPSS 24.0 (IBM, New York, USA) and Fiji (New York, USA). The Shapiro-Wilk test was used to assess the normality of the data and unpaired t-test was used for comparisons between two groups. A two-way analysis of variance (ANOVA) was employed to compare differences among four groups and Tukey’s multiple comparison test was used to compare every two groups. Data are described as the mean ± the standard error of the mean (SEM), and *p* < 0.05 was considered statistically significant.

## Results

### 1. Acute sleep deprivation impaired fear memory

To primarily assess the effect of acute sleep deprivation on cued fear memory, mice underwent 6 hours of sleep deprivation followed by three CS-US pairing trials for fear conditioning training on the first day. After 24h, the expression of fear memory was measured using fear conditioning test across different groups (Fig. 1A). Context control groups were included to evaluate the impact of acute sleep deprivation, which had no impact on fear acquisition, as shown in Fig. 1B. There was no significant difference between Context control group and ASD context control group (Fig. 1C), indicating that acute sleep deprivation had minimal impact on the mice’s response to the context environment. However, ASD-treated mice demonstrated significant impairment of fear memory (Two-way ANOVA test, Turkey’s test analysis: Ctr FC vs ASD FC in CS: *p* = 0.018).

**Figure 1.**
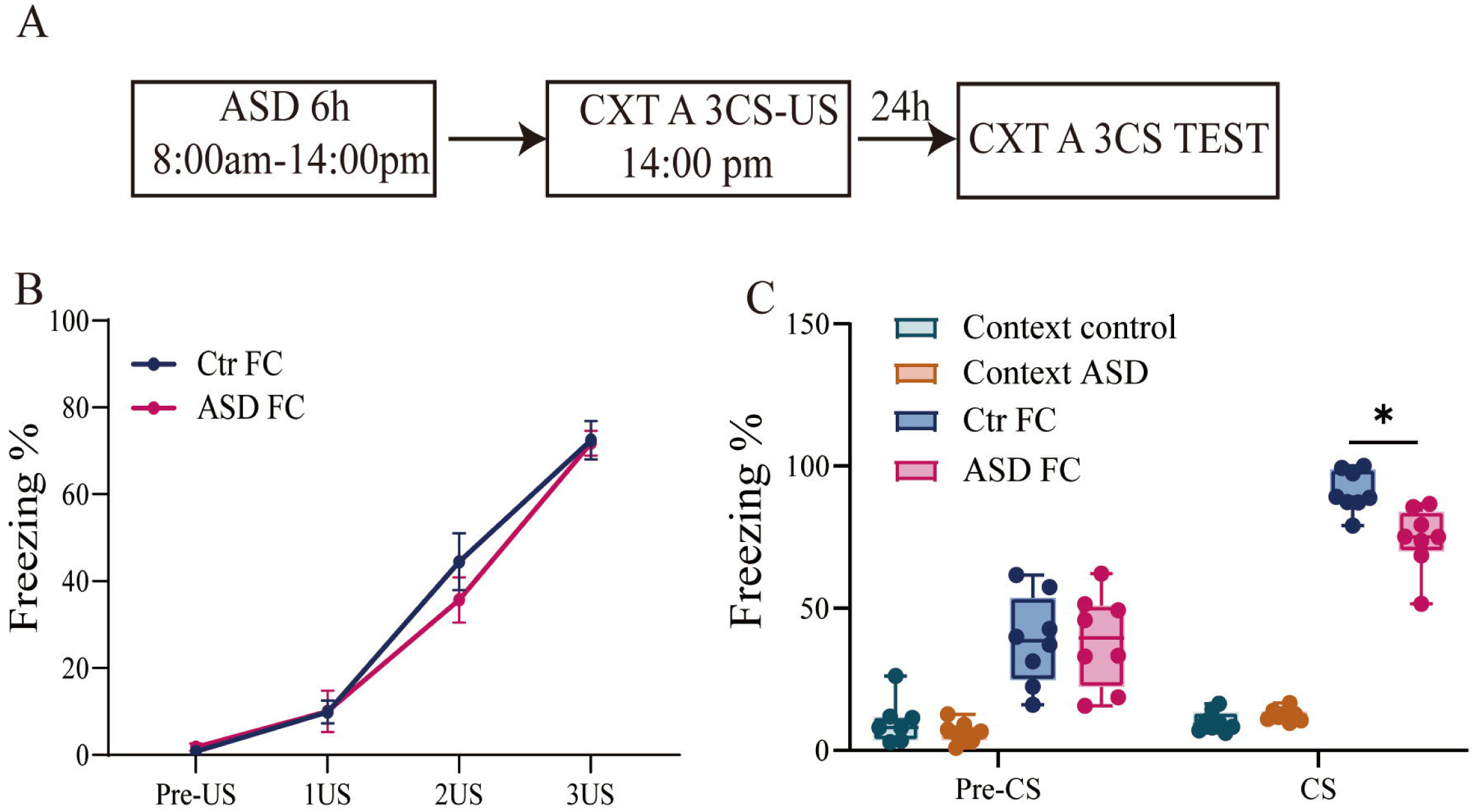
Acute sleep deprivation impaired cued fear memory. A. Time course of acute sleep deprivation and behavior test. B. No significant difference was observed between control and sleep-deprivation groups during fear acquisition. C. Acute sleep deprivation had no effect on the percentage of freezing in the context control and context ASD groups. Sleep-deprived mice showed a significant reduce in the percentage freezing (n=8 per group). FC: Fear conditioned; ASD: acute sleep deprivation; Median ± SEM. \**p* < 0.05.

### 2. mRNA sequence revealed alterations of different genes in fear memory impaired mice induced by sleep deprivation

After sleep-deprived mice underwent fear conditioning training, mPFC tissue was collected for RNA sequencing (Fig. 2A). We compared the transcriptional profile alterations between sleep-deprived mice and those allowed to sleep ad libitum after fear conditioning training. Fig. 2B-C demonstrates the differentially expressed genes (DEGs) in the ASD and Ctr groups. Compared to the Ctr group, the ASD group exhibited 99 downregulated and 115 upregulated genes. Furthermore, GO analysis was performed to identify enriched biological processes and molecular functions in the two groups (Fig. 2D). Some DEGs upregulated in sleep-deprived mice were associated with “apoptotic process”, “plasma membrane” and “calmodulin binding”. In contrast, the downregulated DEGs were enriched for “adenylate cyclase activity” and “mitotic spindle”. Several genes were selected to verify the reliability of mRNA sequence and RT-qPCR was used to verify the mRNA expression in both groups (Fig. 2E-G, n=6, unpaired *t* test). Fig. 2E illustrated the correlation between the expression levels of some genes from DEG analysis (X-axis) and from RT-qPCR analysis (Y-axis) in two groups. In ASD FC group, the downregulation of genes (Adnp, Dbp, Henmt1) identified through DEG analysis was confirmed by RT-qPCR and the upregulated genes (Pim1. Htr5b and Ntpx2) were similarly confirmed using RT-qPCR, indicating the DEG analysis from mRNA sequence is reliable and stable. Compared to the Ctr group, the mRNA expression of Pim1, Htr5b and Ntpx2 was significantly increased in the ASD group (all *p* < 0.01 in Fig. 2F). In addition, the mRNA expression of Adnp (*p* = 0.013), Dbp (*p* < 0.01), and Henmt1 (*p* < 0.01) was significantly downregulated in the ASD group (Fig. 2G).

**Figure 2.**
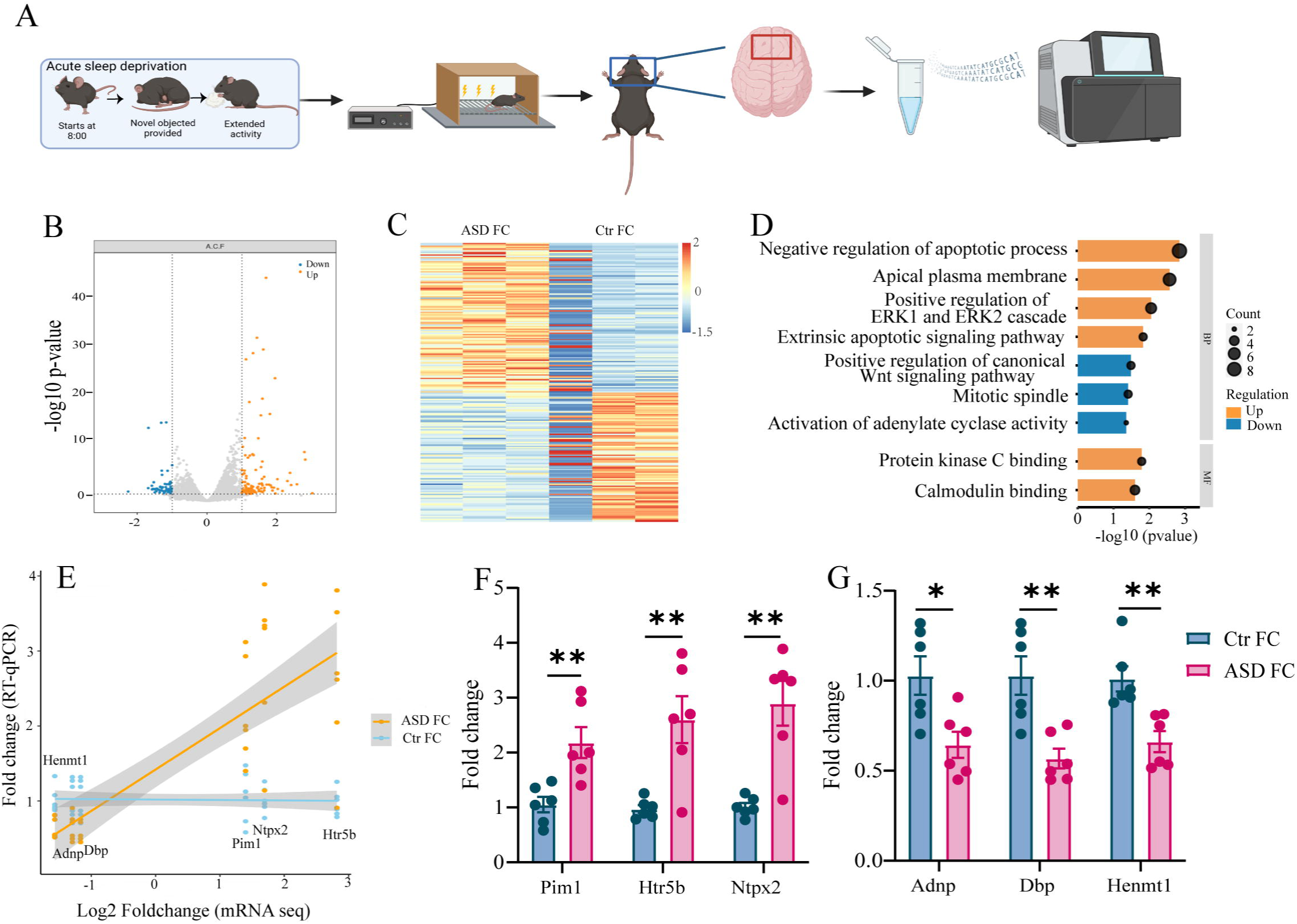
Characterization of Differentially expressed genes (DEGs) between the control and ASD groups. A. Time course of the behavioral modeling and mRNA sequencing. B. Volcano plots representing the upregulated (yellow) and downregulated (blue) DEGs in ASD vs. Ctr in the PL. C. Heatmap of DEG using data from mRNA sequence in two groups (*p* < 0.05). D. Gene Ontology (GO) term analysis of ASD FC vs. Ctr FC, n=3 per group, *p* value < 0.05. E. The correlation between the expression levels of some genes from DEG analysis (X-axis) and RT-qPCR analysis (Y-axis) in two groups. F-G. Verification of upregulated genes and downregulated genes via RT-qPCR in two groups. FC: Fear conditioned; ASD: acute sleep deprivation; Median ± SEM. \**p* < 0.05, \*\**p* < 0.01.

### 3. Enhanced expression of TRPV4 were identified in sleep deprivation mice

From GO analysis, we found that TRPV4 is involved in several biological processes and molecular functions enriched in the ASD group, such as “plasma membrane” and “calmodulin binding.” Furthermore, previous studies have demonstrated a close relationship between TRPV4 and learning and memory [49, 50]. Thus, we preliminarily identified TRPV4 as a critical target in fear memory impairment induced by acute sleep deprivation. To investigate the factors affecting TRPV4 alteration, the mRNA level of TRPV4 was measured in context control, context ASD, control FC and ASD FC groups. The results indicated that acute sleep deprivation significantly promoted TRPV4 expression, whereas fear training had no impact on TRPV4 expression. As shown in Fig 3A, there were significant increases in TRPV4 mRNA level (Two-way ANOVA, Bonferroni’s test, context Ctr vs. context ASD, *p* < 0.0001; Ctr FC vs. ASD FC, *p* < 0.0001). Additionally, the expression of TRPV4 proteins was detected in the Ctr FC and ASD FC groups. Results illustrated a significant increase in TRPV4 protein expression in the PL of the ASD FC group (unpaired *t* test, *p* < 0.001, Fig. 3B).

**Figure 3.**
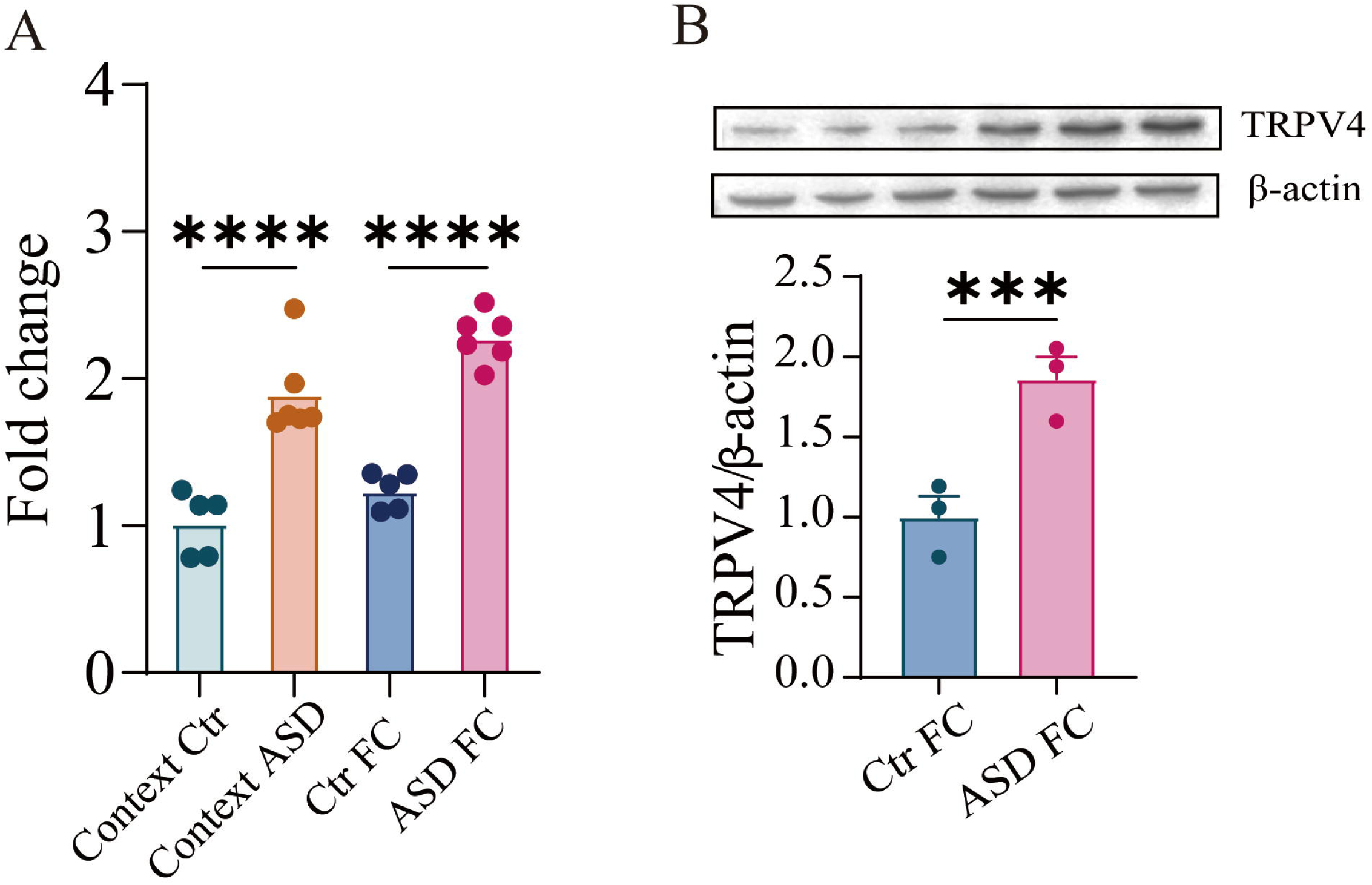
TRPV4 expression is enhanced in the PL after acute sleep deprivation but is not affected by fear condition. A. Acute sleep deprivation led to a significant increase in TRPV4 mRNA expression in both the context control and fear condition groups. n=5-6 per group. B. The expression of TRPV4 protein was significantly enhanced in the ASD FC group compared to Ctr FC group. n=3 per group FC: Fear conditioned; ASD: acute sleep deprivation; Median ± SEM; \**p* < 0.05, \*\*\*\**p* < 0.001.

### 4. The impaired fear memory was associated with enhanced TRPV4 expression caused by acute sleep deprivation

We designed and tested TRPV4-shRNA based on established protocols [22], Subsequently, we packaged shRNA lentivirus targeting TRPV4. The packaged lentivirus was infused into the bilateral PL 7 days before ASD and fear memory behavioral training (Fig. 4A-C). As shown in Fig. 4B, the knockdown efficiency of the TRPV4-shRNA virus was approximately above 50%. Knockdown of TRPV4 did not affect fear memory acquisition (Fig. 4D). However, the percentage of freezing level was significantly elevated in mice infused with the TRPV4-shRNA virus compared to the control virus group after sleep deprivation, indicating that knocking down TRPV4 resulting in the recovery of fear memory in sleep-deprived mice (Fig. 4E, *p =* 0.032). To further investigate how and when TRPV4 influence fear memory, the additional experimental time course was depicted in Fig. 4F. Lentivirus-TRPV4-shRNA or control virus was infused after fear condition training, followed by a behavior test performed 7 days later. As illustrated in Fig. 4G, sleep deprivation did not affect the acquisition of fear memory. The freezing level showed a significant increase in animals infused with TRPV4-shRNA compared to those with the control virus in Fig. 4H, confirming the influence of TRPV4 on fear memory (Two-way ANOVA test, Turkey’s test analysis: *p =* 0.013). All these findings indicate that inhibition of TRPV4 before or after ASD could rescue the fear memory impairment caused by ASD. As TPRV4 is a cation channel related to calcium activity, we further investigated that whether acute sleep deprivation affect the concentration of Ca^2+^ in the PL.

**Figure 4.**
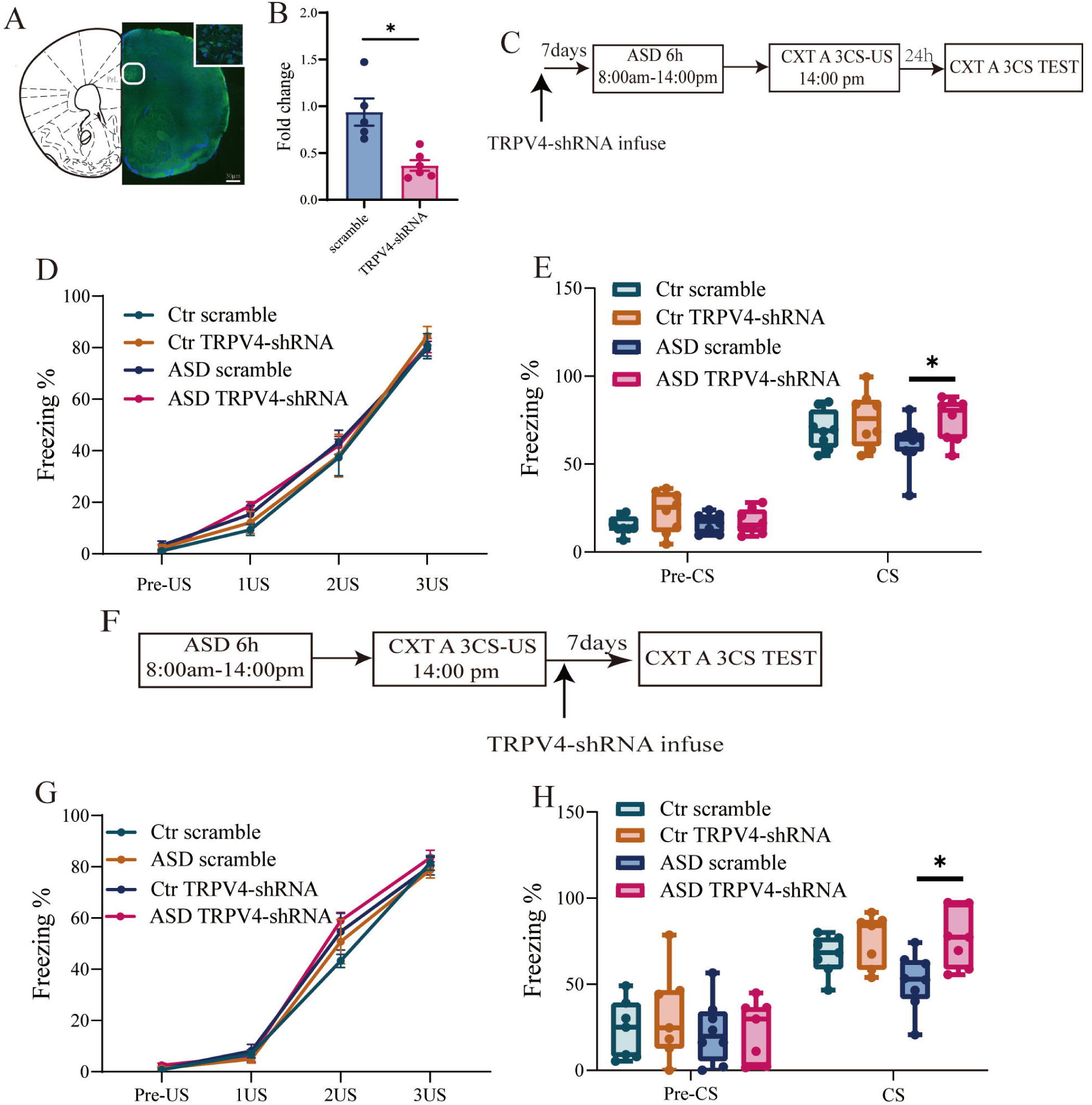
Knocking down TRPV4 in the PL reversed fear memory impairment induced by sleep deprivation. A. Fluorescent images illustrated the location of the TRPV4-shRNA lentivirus infusion. B. The knockdown efficiency of TRPV4-shRNA is approximately 50%. C. Time course of lentivirus infusion, sleep deprivation, and behavioral test. D. TRPV4-shRNA lentivirus had no effect on the acquisition of fear memory. E. The percentage of freezing was significantly increased in animals infused with TRPV4-shRNA in comparison to those infused with control virus after sleep deprivation. F. Time course of sleep deprivation, behavioral training, and lentivirus infusion. G. Sleep deprivation had no effect on fear memory acquisition. H. The freezing level showed a significant increase in ASD animals infused with TRPV4-shRNA compared to control virus. FC: Fear conditioned; ASD: acute sleep deprivation; Median ± SEM. \**p* < 0.05.

### 5. Ca^2+^ overloading and synaptic plasticity damage in mice with fear memory impairment induced by acute sleep deprivation

To explore how sleep deprivation affects fear memory, we examined the concentration of Ca^2+^ and synaptic plasticity change by assessing the expression of PSD95 and the spine density. Schematic of the tissue collection protocols are illustrated in Fig. 5A. ASD significantly increased the concentration of Ca^2+^ in the PL (Fig. 5B, unpaired *t* test, *p* = 0.003). We next investigated the expression of PSD95 protein in mice with fear memory impairment caused by sleep deprivation, as PSD95 is a crucial regulator of synaptic plasticity and is associated with learning and memory. Fig 5C reveals that the mRNA level of PSD95 was significantly decreased (Unpaired *t* test, *p* = 0.025). PSD95 protein expression was also significantly reduced, as depicted in Fig 5D-E (Unpaired *t* test, *p* = 0.019). Representative images from immunofluorescence were depicted in Fig 5F. Quantitative analysis highlighted that the expression of PSD95 protein in the PL was significantly downregulated (Fig. 2G, *p* = 0.0014, unpaired *t* test). Furthermore, Fig. 5H demonstrated time line of tissue collection protocol to examine dendritic spines. Dendritic segments of the PL neurons are shown in Fig. 5I. Segments that can be traced directly to the soma were selected and analyzed. Representative images from the control and ASD group are illustrated in Fig. 5J. The ASD group demonstrated significantly reduced spine density (Unpaired *t* test, Fig. 5K, *p* < .0001).

**Figure 5.**
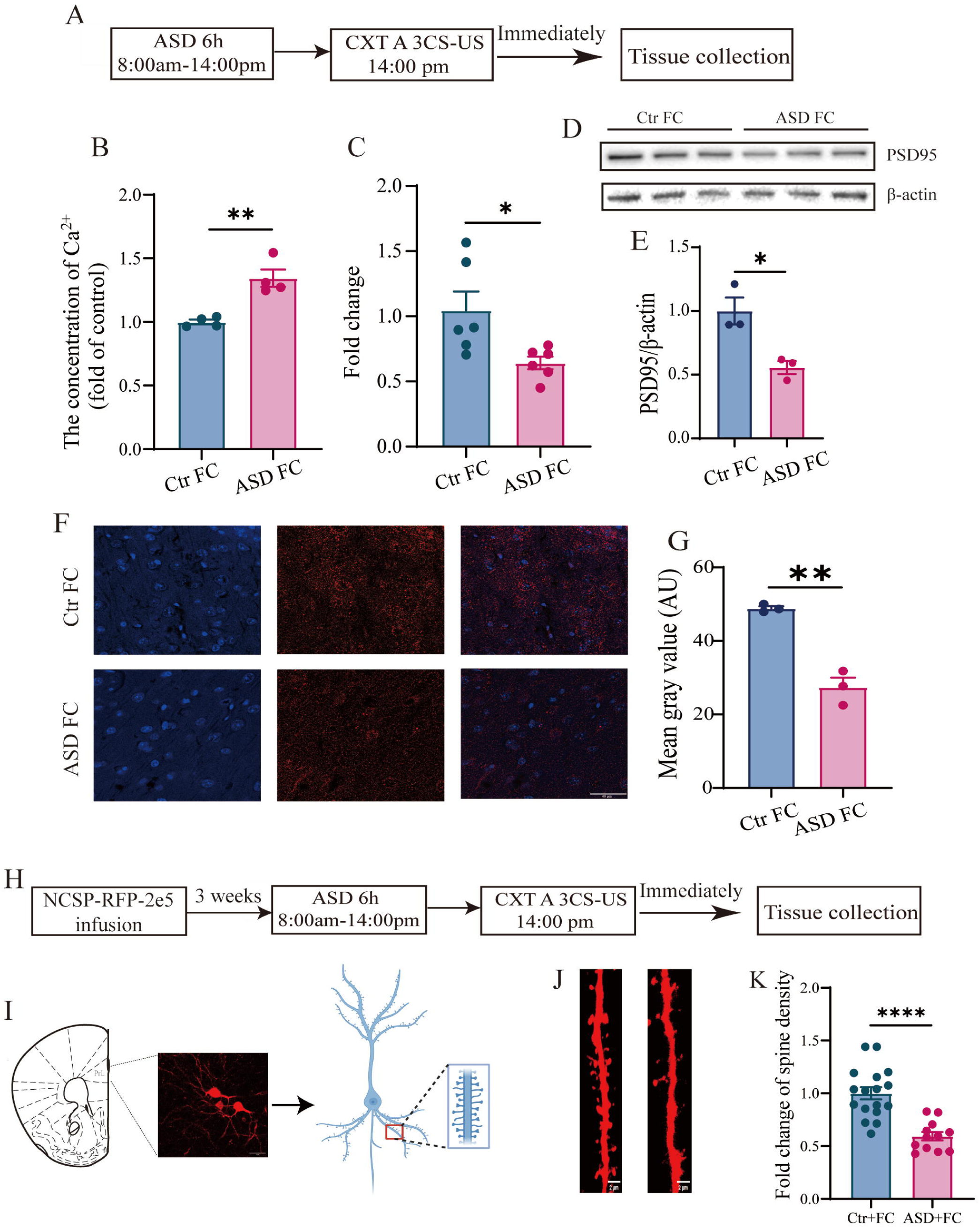
Acute sleep deprivation led to Ca^2+^ overloading and synaptic plasticity impairment in the PL in mice with fear memory impairment. A. Time line of tissue collection for examining the concentration of Ca^2+^ and PSD95 expression. B. The concentration of Ca^2+^ in the PL was significantly increased in ASD FC group. C. The PSD95 mRNA level was significantly decreased in ASD FC group. D. Representative images of immunoblotting of PSD95. E. The expression of PSD95 proteins was significantly reduced in sleep deprived mice. F. Representative images of Immunofluorescence of PSD95 in two groups. G. Quantitative analysis illustrated that the expression of PSD95 proteins was significantly decreased. H. Time line for dendrite spine density analysis. I. Schematic diagram illustrating the dendrite spines of PL neurons. The secondary basal spine dendrites were analyzed. Scale bar: 10µm J. Representative dendrites spines of PL neurons from control and sleep deprived mice. Scale bar: 2µm. K. The spine densities of PL neurons in control and sleep deprived mice were quantified. n=12-15 dendrites from 3 mice per group. FC: Fear conditioned; ASD: acute sleep deprivation; Median ± SEM. \**p* < 0.05, \*\**p* < 0.01, \*\*\**p* < 0.001. Scale bar: 40µm.

### 6. The effect of TRPV4 expression on Ca^2+^ concentration and synaptic plasticity in mice with fear memory impairment after sleep deprivation

Based on above experimental observations, we hypothesized that TRPV4 knockdown alleviates Ca^2+^ overloading and affects synaptic plasticity, thereby improving fear memory. As depicted in Fig. 6A, lentivirus-TRPV4-shRNA was initially infused into the bilateral PL 7 days prior to the experimental day. On the experimental day, mice underwent 6h acute sleep deprivation and were subsequently trained for fear conditioning. Immediately after training, the mice were sacrificed, and their brain tissues were collected for further study. Fig. 6B shows the concentration of Ca2+ was significantly decreased following the infusion of LV-TRPV4-shRNA (Unpaired t test, *p* = 0.012). To elucidate the effect of TRPV4 on synaptic plasticity, we measured the expression of PSD95 protein and dendritic spines alterations and under TRPV4 knocking down conditions. Fig. 6C-E demonstrates that both the mRNA level of PSD95 (Unpaired *t* test, *p* = 0.019) and PSD95 protein (Unpaired *t* test, *p* = 0.005) were significantly increased. Representing immunofluorescent images of PSD95 protein in the PL are showed in Fig. 6F. Quantitative analysis demonstrated a significant increase in PSD95 protein expression following the infusion of LV-TRPV4-shRNA (Unpaired *t* test, *p* = 0.005, Fig. 6G). The time line of tissue collection protocol to examine dendritic spines is shown in Fig. 6H. We selected the dendritic segments which is traced directly to the soma as Fig. 6I demonstrated. Representative fluorescent images from ASD scramble and ASD TRPV4-shRNA are illustrated in Fig. 6J. TRPV4 knockdown reversed the reduced spine density caused by ASD (Unpaired *t* test, Fig. 6K, *p* < .0001).

**Figure 6.**
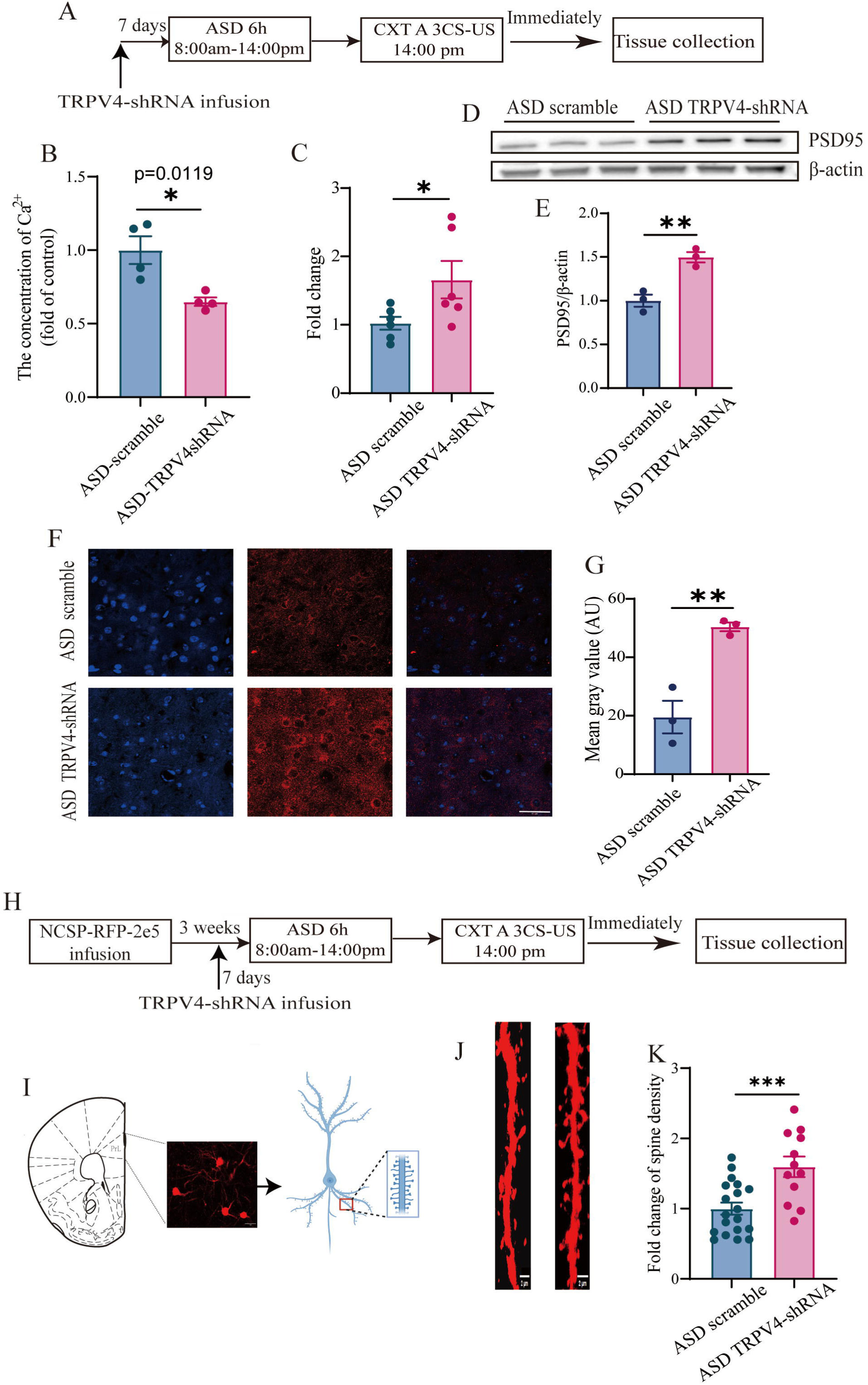
The concentration of Ca^2+^ was reduced and the impaired synaptic plasticity was reversed in the PL after TRPV4-shRNA lentivirus infusion in sleep deprived mice. A. Time line of tissue collection for examining the concentration of Ca^2+^ and PSD95 expression after TRPV4-shRNA knockdown. B. TRPV4 knockdown significantly reduced the concentration of Ca^2+^ in the PL in sleep deprived mice. C. The PSD95 mRNA level was significantly increased in ASD TRPV4-shRNA group. D. Representative images of immunoblotting of PSD95 in two groups. E. The expression of PSD95 proteins was significantly increased after TRPV4-shRNA infusion compared to scramble virus infusion. F. Representative images of Immunofluorescence of PSD95 in two groups. G. Quantitative analysis illustrated that the expression of PSD95 proteins was significantly enhanced. H. Time line for dendrite spine density analysis in different groups. I. Schematic diagram illustrating the dendrite spines of PL neurons. The secondary basal spine dendrites were analyzed. Scale bar: 10µm J. Representative dendrites spines of PL neurons from sleep deprived mice after scramble or TRPV4-shRNA lentivirus infusion. Scale bar: 2µm. K. The spine densities of PL neurons in sleep deprived mice after different virus infusion were quantified. n=12-18 dendrites from 3 mice per group. FC: Fear conditioned; ASD: acute sleep deprivation; Median ± SEM. \**p* < 0.05, \*\**p* < 0.01, \*\*\**p* < 0.001. Scale bar: 40µm.

**Figure 7.**
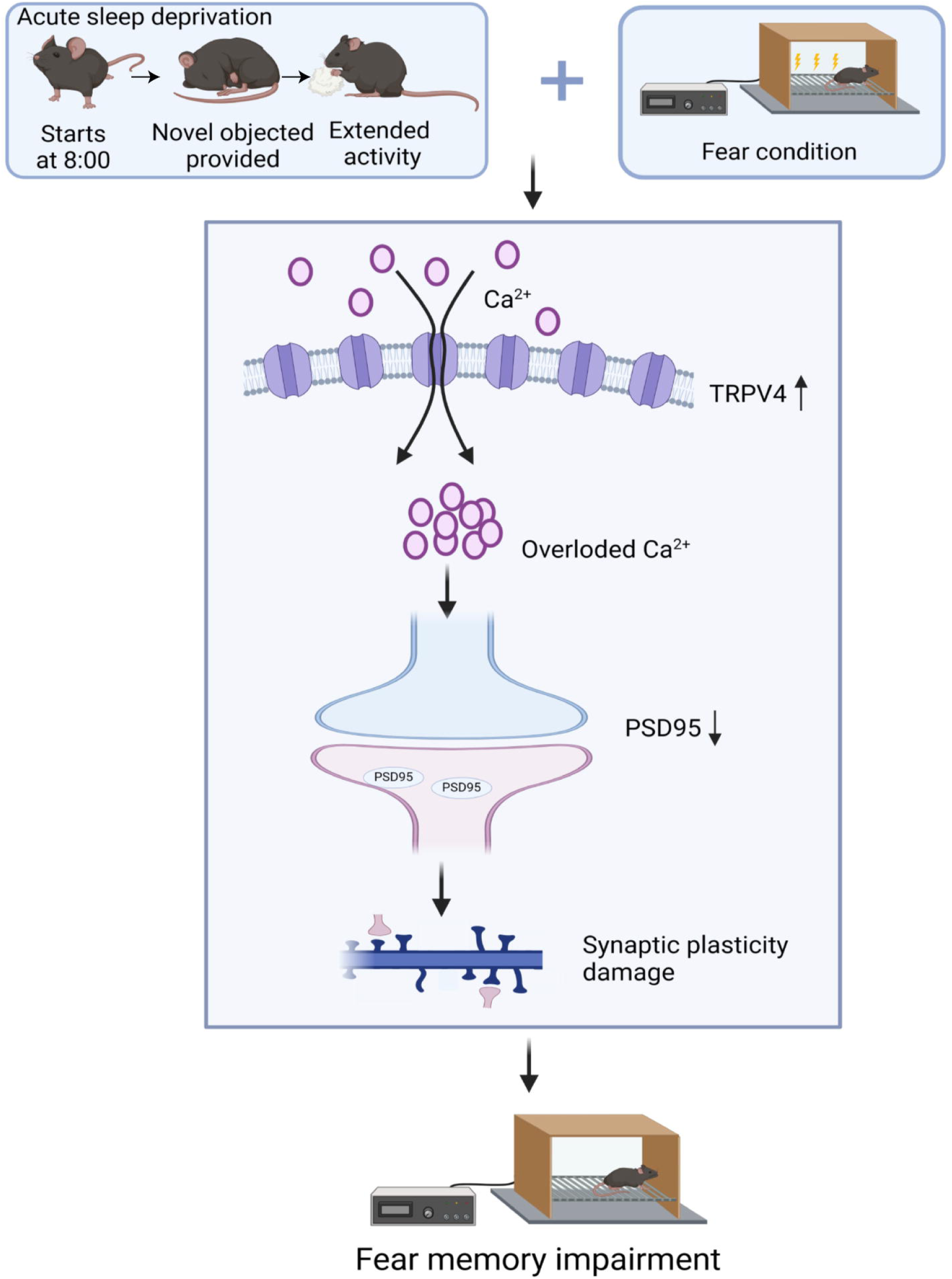
Acute sleep deprivation results in TRPV4 upregulation and Ca2+ overloading in the prelimbic cortex, disrupting synaptic plasticity and impairing fear memory.

## Discussions

In the present study, we explored the effects of sleep deprivation on fear memory. Through mRNA sequence analysis, we identified TRPV4 as a potential target involved in sleep deprivation-induced fear memory impairment. To further investigate the role of TRPV4 in this process, we packaged a lentivirus to specifically reduce TRPV4 expression in the PL. Following lentivirus injection, knockdown of TRPV4 in the PL effectively reversed the fear memory impairment caused by sleep deprivation, concurrently augmenting the loss of dendritic spines and PSD95 protein reduction associated with sleep deprivation. This implied the increase in TRPV4 due to sleep deprivation leads to dendritic spine damage and PSD95 protein reduction, ultimately resulting in fear memory impairment. Altogether, these findings highlight the significant role of TRPV4 in sleep deprivation-induced fear memory impairment and its potential impact on synaptic plasticity. This study underscores the potential of targeting TRPV4 as a therapeutic approach for mitigating the adverse cognitive effects of sleep deprivation.

Sleep plays a crucial role in memory consolidation [49, 50]. Accumulating evidences have demonstrated that neuronal networks show rhythmic activities [51]. Given the important role of neuronal activities in synaptic plasticity, sleep regulates synaptic connections which are necessary for memory learning and formation [52, 53]. Nevertheless, how sleep deprivation affects the memory formation. This study found that acute sleep deprivation impaired synaptic morphological and functional plasticity, leading to impaired fear memory. Previous study also found 8h sleep deprivation significantly reduced spine formation and affect learning and memory. Meanwhile, the reduced spine formation caused by sleep deprivation could not be reversed by additional training [54]. Interestingly, increased dendritic spine density and spin genesis of pyramidal neurons were found 24h after sleep deprivation [55]. Thus, it is a suspect that acute sleep deprivation may lead to reduced spine density and affect learning and memory at that point but the impaired synaptic plasticity might be rescued by subsequent sleep and rest.

Neural activities in the brain are intricate and highly susceptible to variations in sleep conditions. Sleep deprivation can disturb normal neural transmission, leading to deficits in attention and memory [56]. The prelimbic cortex (PL), a subregion of the median prefrontal cortex (mPFC), not only regulates the sleep-wake cycle but also plays a critical role in cognitive processes [57]. Previous studies have shown that sleep disturbances and fragment often precede cognitive deficits in alcohol-dependent patients with abnormal mPFC function [58]. Furthermore, chronic sleep deprivation has been found to suppress neural networks related to the PL and impair synaptic plasticity [59], illustrating that sleep deprivation can induce cognitive dysfunction through abnormal neural activity in the PL. In addition, the PL is also a crucial region for fear memory retrieval [60]. Activation of PL neurons has been associated with increased freezing behavior during fear expression [23] while inactivation of these neurons suppressed fear learning [61, 62]. Given these findings, it is essential to elucidate the regulatory mechanism within the prelimbic cortex that contribute to fear memory impairment caused by acute sleep deprivation.

In the present study, we found that 6h acute sleep deprivation enhances the expression of TRPV4. TRPV4 appears to be particularly sensitive to sleep loss, with its expression pattern is associated with the circadian rhythms of clock genes [28, 63]. TRPV4 levels were elevated during wakefulness and decreased during sleep stages [28]. Interestingly, previous research has shown that members of the TRPV family are involved in regulating sleep homeostasis. For instance, TRPV1, another member of the TRPV family, plays a crucial role in sleep regulation [64], and has been implicated in sleep deprivation-induced fear behavior damage and memory impairment [65]. Additionally, studies have highlighted the significant role of TRPV channels in the sleep-wake cycle. One kind of mechanosensory neurons that express TRPV channels can indirectly affect the sleep center in brain and regulate circadian rhythms [66]. Therefore, the expression of TRPV family, including TRPV4, can mediate sleep phases which in turn can be regulated by sleep deprivation.

TRPV4 channels are novel plasma membrane Ca^2+^ channels that are widely distributed in the brain [67]. As the Ca^2+^ channels are vital for neuronal transmission, alterations of TRPV4 channel might contribute to neurobiological diseases, including Parkinson’s disease [68], Alzheimer’s disease [67], brain edema [69] and others. Previous study reported that TRPV4 inhibitor HC067047 alleviated the learning and memory impairment caused by scopolamine, indicating the promising neuroprotective impact of suppressing TRPV4 [70]. TRPV4 activation triggers the influx of Ca^2+^ from the extracellular matrix and intracellular Ca^2+^ release, leading to an elevation in overall Ca^2+^ concentration [71, 72]. Ca^2+^ serves as a vital messenger contributing to various cellular activities. Imbalances in Ca^2+^ homeostasis is closely associated with neurological diseases. In current study, the expression of TRPV4 was enhanced in ASD-induced fear memory impairment. Notably, knocking down TRPV4 in the PL reversed the fear memory impairment caused by sleep deprivation and led to an increase in spine densities and PSD95 protein levels. These implies that hyperactivation of TRPV4 due to sleep deprivation might lead to Ca^2+^ overload and subsequently impair synaptic plasticity, resulting in fear memory deficits. Similarly, previous study illustrated that knocking out another Ca^2+^-permeable channel TRPC6 improved the balance of Ca^2+^ homeostasis, alleviated neuronal damage and mitigated cognitive deficits [43]. Moreover, Ca^2+^ overload caused by the hyperactivation of TRPV4 has been linked to neuronal inflammation and apoptosis in Parkinson’s disease [68].

### Conclusions

In summary, acute sleep deprivation leads to fear memory impairment with an increase of TRPV4 expression and Ca^2+^ overloading. Additionally, knocking down TRPV4 alleviated synaptic plasticity damage, reversing fear memory deficits in sleep deprived mice. This study suggests that TRPV4 might be a promising therapeutic target for mitigating cognitive impairments associated with sleep deprivation and highlights the importance of sleep management in PTSD patients.

## Acknowledgements

This work was supported by the National Natural Science Foundation of China (82001421 and 82171517). The graphical abstract was created with biorender.com.

## Author contributions

MM Guo, FY Zhang and S Liu collected data. MM Guo and LS Wang performed data analysis. MM Guo, Y Zhang and W Wei drafted the manuscript. J Song, W Wei and X Li conceived the study and finalized the manuscript.

## Data Availability

The data that support the findings of current study are available from the corresponding authors upon reasonable request

## References

1. Puentes-Mestril C, Aton SJ (2017) Linking Network Activity to Synaptic Plasticity during Sleep: Hypotheses and Recent Data. Frontiers in neural circuits 11:61. 10.3389/fncir.2017.00061

2. Rasch B, Born J (2013) About sleep’s role in memory. Physiological reviews 93:681–766. 10.1152/physrev.00032.2012

3. Diekelmann S, Born J (2010) The memory function of sleep. Nature reviews Neuroscience 11:114–26. 10.1038/nrn2762

4. Pace-Schott EF, Germain A, Milad MR (2015) Effects of sleep on memory for conditioned fear and fear extinction. Psychological bulletin 141:835–57. 10.1037/bul0000014

5. Ognjanovski N, Broussard C, Zochowski M, Aton SJ (2018) Hippocampal Network Oscillations Rescue Memory Consolidation Deficits Caused by Sleep Loss. Cerebral cortex (New York, NY: 1991) 28:3711-23. 10.1093/cercor/bhy174

6. Tiba PA, Oliveira MG, Rossi VC, Tufik S, Suchecki D (2008) Glucocorticoids are not responsible for paradoxical sleep deprivation-induced memory impairments. Sleep 31:505–15. 10.1093/sleep/31.4.505

7. Havekes R, Vecsey CG, Abel T (2012) The impact of sleep deprivation on neuronal and glial signaling pathways important for memory and synaptic plasticity. Cellular signalling 24:1251–60. 10.1016/j.cellsig.2012.02.010

8. Gao H, Zhang Y, Luo D, Xu J, Tan S, Li Y, et al. (2023) Activation of the Hippocampal DRD2 Alleviates Neuroinflammation, Synaptic Plasticity Damage and Cognitive Impairment After Sleep Deprivation. Molecular neurobiology 60:7208–21. 10.1007/s12035-023-03514-5

9. Ravassard P, Hamieh AM, Joseph MA, Fraize N, Libourel PA, Lebarillier L, et al. (2016) REM Sleep-Dependent Bidirectional Regulation of Hippocampal-Based Emotional Memory and LTP. Cerebral cortex (New York, NY: 1991) 26:1488-500. 10.1093/cercor/bhu310

10. Kuriyama K, Soshi T, Kim Y (2010) Sleep deprivation facilitates extinction of implicit fear generalization and physiological response to fear. Biological psychiatry 68:991–8. 10.1016/j.biopsych.2010.08.015

11. Shalev A, Liberzon I, Marmar C (2017) Post-Traumatic Stress Disorder. The New England journal of medicine 376:2459–69. 10.1056/NEJMra1612499

12. Dong Y, Li S, Lu Y, Li X, Liao Y, Peng Z, et al. (2020) Stress-induced NLRP3 inflammasome activation negatively regulates fear memory in mice. Journal of neuroinflammation 17:205. 10.1186/s12974-020-01842-0

13. Luchkina NV, Bolshakov VY (2019) Mechanisms of fear learning and extinction: synaptic plasticity-fear memory connection. Psychopharmacology 236:163–82. 10.1007/s00213-018-5104-4

14. Lesuis SL, Lucassen PJ, Krugers HJ (2019) Early life stress impairs fear memory and synaptic plasticity; a potential role for GluN2B. Neuropharmacology 149:195–203. 10.1016/j.neuropharm.2019.01.010

15. Santos TB, Kramer-Soares JC, Favaro VM, Oliveira MGM (2017) Involvement of the prelimbic cortex in contextual fear conditioning with temporal and spatial discontinuity. Neurobiology of learning and memory 144:1–10. 10.1016/j.nlm.2017.05.003

16. Klavir O, Prigge M, Sarel A, Paz R, Yizhar O (2017) Manipulating fear associations via optogenetic modulation of amygdala inputs to prefrontal cortex. Nature neuroscience 20:836–44. 10.1038/nn.4523

17. Widagdo J, Zhao QY, Kempen MJ, Tan MC, Ratnu VS, Wei W, et al. (2016) Experience-Dependent Accumulation of N6-Methyladenosine in the Prefrontal Cortex Is Associated with Memory Processes in Mice. The Journal of neuroscience: the official journal of the Society for Neuroscience 36:6771–7. 10.1523/jneurosci.4053-15.2016

18. Dixsaut L, Gräff J (2022) Brain-wide screen of prelimbic cortex inputs reveals a functional shift during early fear memory consolidation. eLife 11. 10.7554/eLife.78542

19. Rizzo V, Touzani K, Raveendra BL, Swarnkar S, Lora J, Kadakkuzha BM, et al. (2017) Encoding of contextual fear memory requires de novo proteins in the prelimbic cortex. Biological psychiatry Cognitive neuroscience and neuroimaging 2:158–69. 10.1016/j.bpsc.2016.10.002

20. Sun W, Li X, An L (2018) Distinct roles of prelimbic and infralimbic proBDNF in extinction of conditioned fear. Neuropharmacology 131:11–9. 10.1016/j.neuropharm.2017.12.018

21. Burgos-Robles A, Vidal-Gonzalez I, Quirk GJ (2009) Sustained conditioned responses in prelimbic prefrontal neurons are correlated with fear expression and extinction failure. The Journal of neuroscience: the official journal of the Society for Neuroscience 29:8474–82. 10.1523/jneurosci.0378-09.2009

22. Li X, Marshall PR, Leighton LJ, Zajaczkowski EL, Wang Z, Madugalle SU, et al. (2019) The DNA Repair-Associated Protein Gadd45γ Regulates the Temporal Coding of Immediate Early Gene Expression within the Prelimbic Prefrontal Cortex and Is Required for the Consolidation of Associative Fear Memory. The Journal of neuroscience: the official journal of the Society for Neuroscience 39:970–83. 10.1523/jneurosci.2024-18.2018

23. Vidal-Gonzalez I, Vidal-Gonzalez B, Rauch SL, Quirk GJ (2006) Microstimulation reveals opposing influences of prelimbic and infralimbic cortex on the expression of conditioned fear. Learning & memory (Cold Spring Harbor, NY) 13:728–33. 10.1101/lm.306106

24. Reboreda A, Jiménez-Díaz L, Navarro-López JD (2011) TRP channels and neural persistent activity. Advances in experimental medicine and biology 704:595–613. 10.1007/978-94-007-0265-3_32

25. Kakae M, Nakajima H, Tobori S, Kawashita A, Miyanohara J, Morishima M, et al. (2023) The astrocytic TRPA1 channel mediates an intrinsic protective response to vascular cognitive impairment via LIF production. Science advances 9:eadh0102. 10.1126/sciadv.adh0102

26. Grace MS, Bonvini SJ, Belvisi MG, McIntyre P (2017) Modulation of the TRPV4 ion channel as a therapeutic target for disease. Pharmacology & therapeutics 177:9–22. 10.1016/j.pharmthera.2017.02.019

27. Ihara T, Mitsui T, Nakamura Y, Kanda M, Tsuchiya S, Kira S, et al. (2018) The oscillation of intracellular Ca(2+) influx associated with the circadian expression of Piezo1 and TRPV4 in the bladder urothelium. Scientific reports 8:5699. 10.1038/s41598-018-23115-w

28. Ihara T, Mitsui T, Nakamura Y, Kira S, Nakagomi H, Sawada N, et al. (2017) Clock Genes Regulate the Circadian Expression of Piezo1, TRPV4, Connexin26, and VNUT in an Ex Vivo Mouse Bladder Mucosa. PloS one 12:e0168234. 10.1371/journal.pone.0168234

29. Deng Y, Li W, Niu L, Luo X, Li J, Zhang Y, et al. (2022) Amelioration of scopolamine-induced learning and memory impairment by the TRPV4 inhibitor HC067047 in ICR mice. Neuroscience Letters 767:136209. 10.1016/j.neulet.2021.136209

30. Veteto AB, Peana D, Lambert MD, McDonald KS, Domeier TL (2020) Transient receptor potential vanilloid-4 contributes to stretch-induced hypercontractility and time-dependent dysfunction in the aged heart. Cardiovascular research 116:1887–96. 10.1093/cvr/cvz287

31. Baratchi S, Keov P, Darby WG, Lai A, Khoshmanesh K, Thurgood P, et al. (2019) The TRPV4 Agonist GSK1016790A Regulates the Membrane Expression of TRPV4 Channels. Frontiers in pharmacology 10:6. 10.3389/fphar.2019.00006

32. Bai JZ, Lipski J (2014) Involvement of TRPV4 channels in Aβ(40)-induced hippocampal cell death and astrocytic Ca(2+) signalling. Neurotoxicology 41:64–72. 10.1016/j.neuro.2014.01.001

33. Shi M, Du F, Liu Y, Li L, Cai J, Zhang GF, et al. (2013) Glial cell-expressed mechanosensitive channel TRPV4 mediates infrasound-induced neuronal impairment. Acta neuropathologica 126:725–39. 10.1007/s00401-013-1166-x

34. Lee JC, Choe SY (2014) Age-related changes in the distribution of transient receptor potential vanilloid 4 channel (TRPV4) in the central nervous system of rats. Journal of molecular histology 45:497–505. 10.1007/s10735-014-9578-z

35. Lu KT, Huang TC, Tsai YH, Yang YL (2017) Transient receptor potential vanilloid type 4 channels mediate Na-K-Cl-co-transporter-induced brain edema after traumatic brain injury. Journal of neurochemistry 140:718–27. 10.1111/jnc.13920

36. Jie P, Tian Y, Hong Z, Li L, Zhou L, Chen L, et al. (2015) Blockage of transient receptor potential vanilloid 4 inhibits brain edema in middle cerebral artery occlusion mice. Frontiers in cellular neuroscience 9:141. 10.3389/fncel.2015.00141

37. Jie P, Hong Z, Tian Y, Li Y, Lin L, Zhou L, et al. (2015) Activation of transient receptor potential vanilloid 4 induces apoptosis in hippocampus through downregulating PI3K/Akt and upregulating p38 MAPK signaling pathways. Cell death & disease 6:e1775. 10.1038/cddis.2015.146

38. Jie P, Lu Z, Hong Z, Li L, Zhou L, Li Y, et al. (2016) Activation of Transient Receptor Potential Vanilloid 4 is Involved in Neuronal Injury in Middle Cerebral Artery Occlusion in Mice. Molecular neurobiology 53:8–17. 10.1007/s12035-014-8992-2

39. Cho KO, Hunt CA, Kennedy MB (1992) The rat brain postsynaptic density fraction contains a homolog of the Drosophila discs-large tumor suppressor protein. Neuron 9:929–42. 10.1016/0896-6273(92)90245-9

40. Coley AA, Gao WJ (2018) PSD95: A synaptic protein implicated in schizophrenia or autism? Progress in neuro-psychopharmacology & biological psychiatry 82:187–94. 10.1016/j.pnpbp.2017.11.016

41. Gilman SR, Iossifov I, Levy D, Ronemus M, Wigler M, Vitkup D (2011) Rare de novo variants associated with autism implicate a large functional network of genes involved in formation and function of synapses. Neuron 70:898–907. 10.1016/j.neuron.2011.05.021

42. Bulovaite E, Qiu Z, Kratschke M, Zgraj A, Fricker DG, Tuck EJ, et al. (2022) A brain atlas of synapse protein lifetime across the mouse lifespan. Neuron 110:4057–73.e8. 10.1016/j.neuron.2022.09.009

43. Kong L, Sun R, Zhou H, Shi Q, Liu Y, Han M, et al. (2023) Trpc6 knockout improves behavioral dysfunction and reduces Aβ production by inhibiting CN-NFAT1 signaling in T2DM mice. Experimental neurology 363:114350. 10.1016/j.expneurol.2023.114350

44. Guo M, Wu Y, Zheng D, Chen L, Xiong B, Wu J, et al. (2022) Preoperative Acute Sleep Deprivation Causes Postoperative Pain Hypersensitivity and Abnormal Cerebral Function. Neuroscience bulletin 38:1491–507. 10.1007/s12264-022-00955-1

45. Wei W, Zhao Q, Wang Z, Liau WS, Basic D, Ren H, et al. (2022) ADRAM is an experience-dependent long noncoding RNA that drives fear extinction through a direct interaction with the chaperone protein 14-3-3. Cell reports 38:110546. 10.1016/j.celrep.2022.110546

46. Guo M, Wang J, Yuan Y, Chen L, He J, Wei W, et al. (2023) Role of adenosine A(2A) receptors in the loss of consciousness induced by propofol anesthesia. Journal of neurochemistry 164:684–99. 10.1111/jnc.15734

47. Lin Q, Wei W, Coelho CM, Li X, Baker-Andresen D, Dudley K, et al. (2011) The brain-specific microRNA miR-128b regulates the formation of fear-extinction memory. Nature neuroscience 14:1115–7. 10.1038/nn.2891

48. Zhang YQ, Lin WP, Huang LP, Zhao B, Zhang CC, Yin DM (2021) Dopamine D2 receptor regulates cortical synaptic pruning in rodents. Nature communications 12:6444. 10.1038/s41467-021-26769-9

49. Maquet P (2001) The role of sleep in learning and memory. Science (New York, NY) 294:1048–52. 10.1126/science.1062856

50. Siegel JM (2005) Clues to the functions of mammalian sleep. Nature 437:1264–71. 10.1038/nature04285

51. Crunelli V, Hughes SW (2010) The slow (<1 Hz) rhythm of non-REM sleep: a dialogue between three cardinal oscillators. Nature neuroscience 13:9–17. 10.1038/nn.2445

52. Lichtman JW, Colman H (2000) Synapse elimination and indelible memory. Neuron 25:269–78. 10.1016/s0896-6273(00)80893-4

53. Yang G, Pan F, Gan WB (2009) Stably maintained dendritic spines are associated with lifelong memories. Nature 462:920–4. 10.1038/nature08577

54. Yang G, Lai CSW, Cichon J, Ma L, Li W, Gan W-B (2014) Sleep promotes branch-specific formation of dendritic spines after learning. 344:1173–8. doi:10.1126/science.1249098

55. Wu M, Zhang X, Feng S, Freda SN, Kumari P, Dumrongprechachan V, et al. (2024) Dopamine pathways mediating affective state transitions after sleep loss. Neuron 112:141–54.e8. 10.1016/j.neuron.2023.10.002

56. Krause AJ, Simon EB, Mander BA, Greer SM, Saletin JM, Goldstein-Piekarski AN, et al. (2017) The sleep-deprived human brain. Nature reviews Neuroscience 18:404–18. 10.1038/nrn.2017.55

57. Chauveau F, Laudereau K, Libourel PA, Gervasoni D, Thomasson J, Poly B, et al. (2014) Ciproxifan improves working memory through increased prefrontal cortex neural activity in sleep-restricted mice. Neuropharmacology 85:349–56. 10.1016/j.neuropharm.2014.04.017

58. Liu J, Cai W, Zhao M, Cai W, Sui F, Hou W, et al. (2019) Reduced resting-state functional connectivity and sleep impairment in abstinent male alcohol-dependent patients. Human brain mapping 40:4941–51. 10.1002/hbm.24749

59. Zhu J, Chen C, Li Z, Liu X, He J, Zhao Z, et al. (2023) Overexpression of Sirt6 ameliorates sleep deprivation induced-cognitive impairment by modulating glutamatergic neuron function. Neural regeneration research 18:2449–58. 10.4103/1673-5374.371370

60. DeNardo LA, Liu CD, Allen WE, Adams EL, Friedmann D, Fu L, et al. (2019) Temporal evolution of cortical ensembles promoting remote memory retrieval. Nature neuroscience 22:460–9. 10.1038/s41593-018-0318-7

61. Sierra-Mercado D, Padilla-Coreano N, Quirk GJ (2011) Dissociable roles of prelimbic and infralimbic cortices, ventral hippocampus, and basolateral amygdala in the expression and extinction of conditioned fear. Neuropsychopharmacology: official publication of the American College of Neuropsychopharmacology 36:529–38. 10.1038/npp.2010.184

62. Corcoran KA, Quirk GJ (2007) Activity in prelimbic cortex is necessary for the expression of learned, but not innate, fears. The Journal of neuroscience: the official journal of the Society for Neuroscience 27:840–4. 10.1523/jneurosci.5327-06.2007

63. Ihara T, Mitsui T, Nakamura Y, Kanda M, Tsuchiya S, Kira S, et al. (2018) The Circadian expression of Piezo1, TRPV4, Connexin26, and VNUT, associated with the expression levels of the clock genes in mouse primary cultured urothelial cells. Neurourology and urodynamics 37:942-51. 10.1002/nau.23400

64. Murillo-Rodríguez E, Di Marzo V, Machado S, Rocha NB, Veras AB, Neto GAM, et al. (2017) Role of N-Arachidonoyl-Serotonin (AA-5-HT) in Sleep-Wake Cycle Architecture, Sleep Homeostasis, and Neurotransmitters Regulation. Frontiers in molecular neuroscience 10:152. 10.3389/fnmol.2017.00152

65. Ozathaley A, Kou Z, Ma Y, Luo D, Chen J, Liu C, et al. (2023) NLRP3 upregulation related to sleep deprivation-induced memory and emotional behavior changes in TRPV1-/-mice. Behavioural Brain Research 440:114255. 10.1016/j.bbr.2022.114255

66. Lone SR, Potdar S, Venkataraman A, Sharma N, Kulkarni R, Rao S, et al. (2021) Mechanosensory Stimulation via Nanchung Expressing Neurons Can Induce Daytime Sleep in Drosophila. The Journal of neuroscience: the official journal of the Society for Neuroscience 41:9403–18. 10.1523/jneurosci.0400-21.2021

67. Liu N, Yan F, Ma Q, Zhao J (2020) Modulation of TRPV4 and BKCa for treatment of brain diseases. Bioorganic & medicinal chemistry 28:115609. 10.1016/j.bmc.2020.115609

68. Liu N, Bai L, Lu Z, Gu R, Zhao D, Yan F, et al. (2022) TRPV4 contributes to ER stress and inflammation: implications for Parkinson’s disease. Journal of neuroinflammation 19:26. 10.1186/s12974-022-02382-5

69. Faropoulos K, Polia A, Tsakona C, Pitaraki E, Moutafidi A, Gatzounis G, et al. (2021) Evaluation of AQP4/TRPV4 Channel Co-expression, Microvessel Density, and its Association with Peritumoral Brain Edema in Intracranial Meningiomas. Journal of molecular neuroscience: MN 71:1786–95. 10.1007/s12031-021-01801-1

70. Deng Y, Li W, Niu L, Luo X, Li J, Zhang Y, et al. (2022) Amelioration of scopolamine-induced learning and memory impairment by the TRPV4 inhibitor HC067047 in ICR mice. Neuroscience letters 767:136209. 10.1016/j.neulet.2021.136209

71. Caterina MJ, Schumacher MA, Tominaga M, Rosen TA, Levine JD, Julius DJN (1997) The capsaicin receptor: a heat-activated ion channel in the pain pathway. 389:816–24.

72. Senning EN, Gordon SEJE (2015) Activity and Ca2+ regulate the mobility of TRPV1 channels in the plasma membrane of sensory neurons. 4:e03819.

